# Polymorphic SNPs, short tandem repeats and structural variants are responsible for differential gene expression across C57BL/6 and C57BL/10 substrains

**DOI:** 10.1101/2020.03.16.993683

**Authors:** Milad Mortazavi, Yangsu Ren, Shubham Saini, Danny Antaki, Celine St. Pierre, April Williams, Abhishek Sohni, Miles Wilkinson, Melissa Gymrek, Jonathan Sebat, Abraham A. Palmer

**Affiliations:** Department of Psychiatry, University of California San Diego, La Jolla, CA; Department of Computer Science and Engineering, University of California San Diego, La Jolla, CA; Department of Cellular and Molecular Medicine and Pediatrics, University of California San Diego, La Jolla, CA; Department of Genetics, Washington University School of Medicine, St. Louis MO; Salk Institute for Biological Studies, La Jolla, CA; Department of Obstetrics, Gynecology and Reproductive Sciences, University of California San Diego, La Jolla, CA; Institute for Genomic Medicine, University of California San Diego, La Jolla, CA; Department of Medicine, University of California San Diego, La Jolla, CA

**Keywords:** C57BL/6, C57BL/10, Reduced Complexity Cross, substrain, mutational processes, gene expression, introgression, *Wdfy1*, *Lpp*, *Srp54*

## Abstract

Mouse substrains are an invaluable model for understanding disease. We compared C57BL/6J, which is the most commonly used inbred mouse strain, with 8 C57BL/6 and 5 C57BL/10 closely related inbred substrains. Whole genome sequencing and RNA-sequencing analysis yielded 352,631 SNPs, 109,096 INDELs, 150,344 short tandem repeats (STRs), 3,425 structural variants (SVs) and 2,826 differentially expressed genes (DEGenes) among these 14 strains. 312,981 SNPs (89%) distinguished the B6 and B10 lineages. These SNPs were clustered into 28 short segments that are likely due to introgressed haplotypes rather than new mutations. Outside of these introgressed regions, we identified 53 SVs, protein-truncating SNPs and frameshifting INDELs that were associated with DEGenes. Our results can be used for both forward and reverse genetic approaches, and illustrate how introgression and mutational processes give rise to differences among these widely used inbred substrains.

## 1. Introduction

Since Clarence C. Little generated the C57BL/6 inbred strain a century ago, the C57BL/6J has become the most commonly used inbred mouse strain. Closely-related C57BL/10 substrains^1,2^, which were separated from C57BL/6 in about 1937, are also commonly used in specific fields such as immunology^3^ and muscular dystrophy^4^. The popularity of inbred C57BL strains has led to the establishment of many substrains (defined as >20 generations of separation from the parent colony). Among the C57BL/6 branches, the two predominant lineages are based on C57BL/6J (from The Jackson Laboratory; **JAX**) and C57BL/6N (from the National Institutes of Health; **NIH**^5,6^). Subsequently, several additional substrains have been derived from the JAX and the NIH branches.

Genetic differences between closely-related laboratory strains have been assumed to be the result of accumulated spontaneous mutations^7^. For those that are selectively neutral, genetic drift dictates that some new mutations will be lost, others will maintain an intermediate frequency, and others will become fixed, replacing the ancestral allele^8^. Because of historical bottlenecks and small breeding populations, fixation of new mutations can be relatively rapid.

Numerous studies have reported phenotypic differences among various C57BL/6- and C57BL/10-derived substrains, which are likely attributable to genetic variation. For C57BL/6 substrains, these differences include learning behavior^9^, prepulse inhibition^10^, anxiety and depression^11^, fear conditioning^12-14^, glucose tolerance^15^, alcohol-related behaviors^16,17^, and responses to other various drugs^18-21^. For C57BL/10 substrains, these differences include seizure traits^22^ and responses to drugs^23^. Crosses between two phenotypically divergent strains can be used for quantitative trait mapping. Because crosses among closely related substrains segregate fewer variants than crosses of more divergent strains, identification of causal alleles is greatly simplified^21^. Such crosses have been referred to as a reduced complexity cross (**RCC**)^24^ and have been further simplified by the recent development of an inexpensive microarray explicitly designed for mapping studies that use RCCs^25^.

Whole Genome Sequencing (**WGS**) technology provides a deep characterization of Single Nucleotide Polymorphisms (**SNPs**), small insertions and deletions (**INDELs**), Short Tandem Repeats (**STRs**), and Structural Variations (**SVs**). SNPs that differentiate a few of the C57BL/6 substrains have been previously reported^21,26^. While most SNPs are expected to have no functional consequences, a subset will; for example, SNPs in regulatory and coding regions, which can profoundly alter gene expression and function. STRs have never been systematically studied in C57BL substrains. STRs are highly variable elements that play a pivotal role in multiple genetic diseases, population genetics applications, and forensic casework. STRs exhibit rapid mutation rates of ∼10^−5^ mutations per locus per generation^27^, orders of magnitude higher than that of point mutations (∼10^−8^)^28^, and are known to play a key role in more than 30 Mendelian disorders^29^. Recent evidence has underscored the profound regulatory role of STRs, suggesting widespread involvement in complex traits^30^. SVs include deletions, duplications, insertions, inversions, and translocations. SVs are individually less abundant than SNPs and STRs, but collectively account for a similar proportion of overall sequence difference between genomes^31^. In addition, SVs can have greater functional consequences because they can result in large changes to protein coding exons or regulatory elements^32^. Large SVs among C57BL/6 (but not C57BL10) substrains were identified using array comparative genomic hybridization^7^, and have also been identified in more diverse panels of inbred strains using WGS^33^. Although some genetic variants that differ between closely related C57BL substrains have been previously reported^7,34-36^, a comprehensive, genome-wide map of SNPs, INDELs, STRs, SVs and gene expression differences among C57BL6 and C57BL10 substrains does not exist.

In an effort to create such a resource, we performed whole genome sequencing in a single male individual from 9 C57BL/6 and 5 C57BL/10 substrains (∼30x per substrain) and called SNPs, INDELs, STRs and SVs. In addition, to identify functional consequences of these polymorphisms, we performed RNA-sequencing of the hippocampal transcriptome in 6-11 male mice from each substrain, which allowed us to identify genes that are differentially expressed (**Figure 1A**). This approach has two advantages: first it provides a large number of molecular phenotypes that may be caused by substrain specific polymorphisms. Second, we assumed that the gene expression differences would often reflect the action of cis regulatory variants, making it possible to narrow the number of potentially causal mutations without requiring the creation of intercrosses.

**Figure 1.**
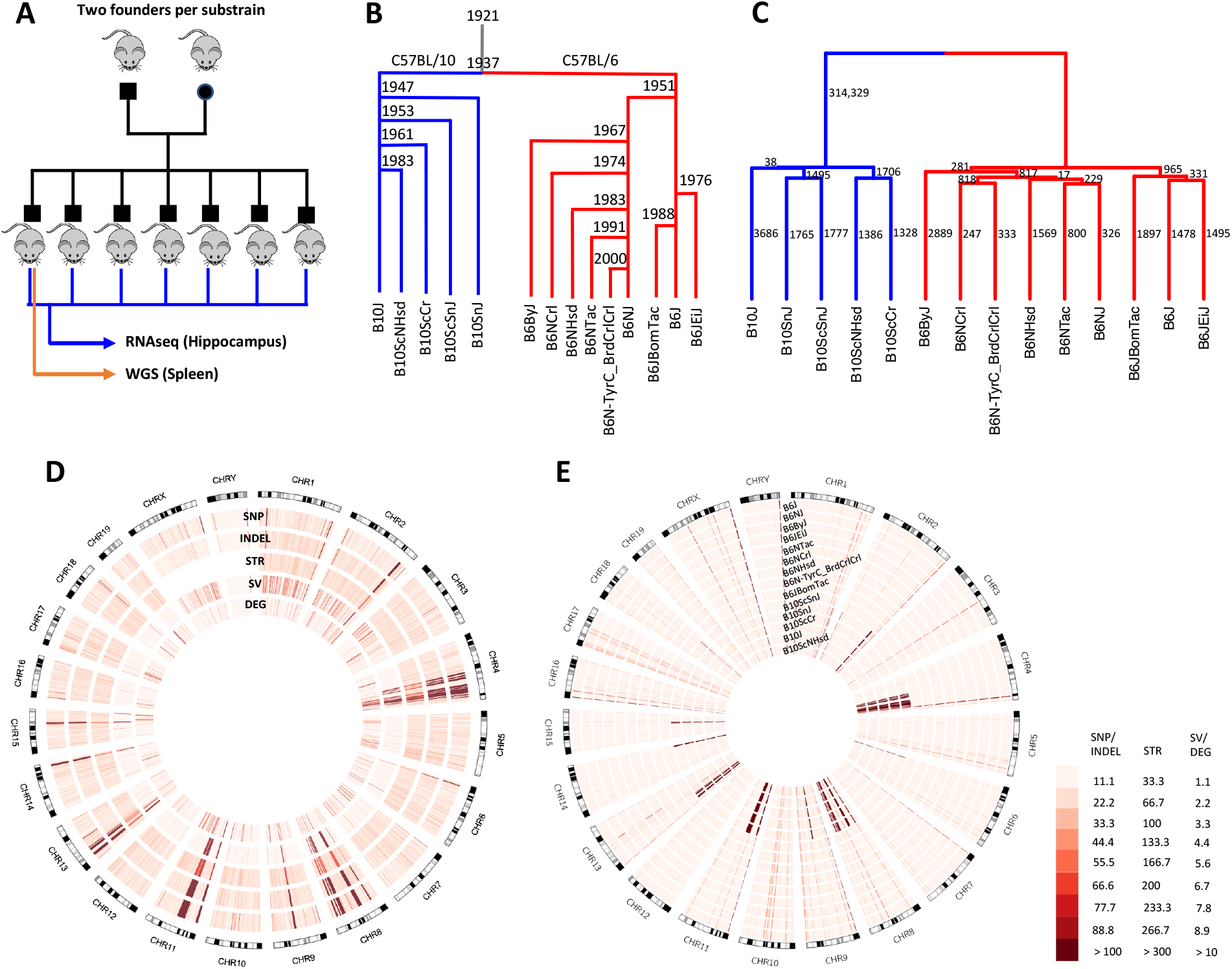
Study design, genetic distance analysis and distribution of genomic variants across the genome. **A**: This panel shows the design of our study. Mice from nine C57BL/6 and five C57BL/10 substrains were purchased from four vendors. For each substrain, between six and eleven male offspring from the first generation born in our colony were chosen for hippocampal RNA-sequencing (RNA-Seq). One male offspring per substrain was chosen for whole genome sequencing (WGS) using DNA extracted from the spleen. **B**: Historical relationship of C57BL/6 and C57BL/10 substrains is illustrated as a tree^37-39^. The year in which each substrain was separated from its branch is shown at its junction. **C**: Dendrogram showing the similarity of the substrains based on genomic variants including: SNPs (LD-pruned, INDELs not included), STRs and SVs. The numbers besides each branch indicates the number of SNPs separating the substrains in the subtree below the branch from all the other substrains. **D**: Circus plot showing the SNPs, INDELs, STRs, SVs, and DEGenes across the genome for 14 C57BL/6 and C57BL/10 substrains. Regions with a high density of polymorphisms (hot spots) on chromosomes 4, 8, 9, 11 and 13 are obvious (see also **Figure S2** and **Figure S3**). **E**: Circus plot showing SNPs with non-reference genotypes for each substrain. This plot shows that most hot spots in panel D are due to regions where all C57BL/6 differ from all C57BL/10 substrains (see also **Figure S2** and **Figure S3**). A few regions where all substrains (including C57BL/6J) do not match the reference are also evident.

## 2. Results

Processing of WGS data from the 14 C57BL substrains (**Table 1**) allowed us to identify 352,631 SNPs, 109,096 INDELs, 150,344 polymorphic STRs and 3,425 SVs in nine C57BL/6 and five C57BL/10 substrains. 5.6% of SNPs and 17.2% of INDELs are singletons (only occur in one substrain). 89% of SNPs and 58% of INDELs separated the C57BL/6 and C57BL/10 branches. The fraction of variants in each category observed in different numbers of substrains is plotted in **Figure S1**.

**Table 1.** Information about samples, WGS and RNA-Seq data used in this study.

| Strain | Vendor | Strain ID | Whole Genome Sequencing |  | RNA-sequencing |  |
| --- | --- | --- | --- | --- | --- | --- |
|  |  |  | Reads | Coverage | Average reads | # samples |
| C57BL/6Ntac | Taconic | B6 | 336,470,599 | 33.65 | 13,459,345 | 8 |
| C57BL/6NJ | JAX | #005304 | 333,094,625 | 33.31 | 15,767,477 | 9 |
| C57BL/6NHsd | Harlan | #044 | 349,305,241 | 34.93 | 14,656,480 | 7 |
| C57BL/6NCrl | Charles River | #027 | 301,990,053 | 30.20 | 19,408,390 | 8 |
| C57BL/6JeiJ | JAX | #000924 | 442,604,628 | 44.26 | 16,334,723 | 8 |
| C57BL/6JbomTac | Taconic | B6Jbom | 309,053,911 | 30.91 | 17,340,684 | 7 |
| C57BL/6J | JAX | #000664 | 326,208,826 | 32.62 | 18,555,108 | 8 |
| C57BL/6ByJ | JAX | #001139 | 294,061,881 | 29.41 | 15,552,591 | 9 |
| B6N-TyrC/BrdCrCrl | Charles River | #493 | 316,655,519 | 31.67 | 15,139,469 | 7 |
| C57BL/10SnJ | JAX | #000666 | 284,191,454 | 28.42 | 14,012,862 | 6 |
| C57BL/10ScSnJ | JAX | #000476 | 285,201,231 | 28.52 | 18,947,235 | 7 |
| C57BL/10ScNHsd | Taconic | #046 | 326,586,425 | 32.66 | 12,453,407 | 11 |
| C57BL/10ScCr | JAX | #003752 | 289,068,061 | 28.91 | 16,384,792 | 8 |
| C57BL/10J | JAX | #000665 | 311,292,991 | 31.13 | 17,550,456 | 7 |
| Average |  |  | 321,841,818 | 32.18 | 16,013,858 | 7.86 |

RNA-sequencing analysis on 106 hippocampal samples identified 16,400 expressed genes and 2,826 DEGenes (17.2%) in C57BL/6 and C57BL/10 substrains (FDR<0.05).

RNA-sequencing data was also used to validate WGS SNPs and INDELs in protein coding regions of the mouse genome (See STAR Methods, **Figure S1** and **Table S3**). We observed a 97% validation rate for SNPs and 67% for INDELs.

Despite the lower sequencing depth on the X chromosome that was expected for male mice, we did not observe a difference in the density of private SNPs discovered on chromosome X compared to the autosomes (7.2 SNPs/Mb on X versus average 7.5 SNPs/Mb on the autosomes). However, the validation rate of coding SNPs on chromosome X was lower than on the autosomes (89% for chromosome X versus 97% on the autosomes).

We also detected 582,795 heterozygous SNP calls. For 92% of these heterozygous calls, all samples show heterozygous genotypes, which suggests that most reflect segmental duplications rather than true heterozygous SNPs. Consistent with this idea, many apparently heterozygous SNPs were in known segmental duplication or tandem repeat regions (45%). For the remaining apparently heterozygous SNPs, we obtained a very low validation rate when using our RNA-Seq data (12% validation rate for heterozygous SNPs in protein coding regions that were not in known segmental duplication or tandem repeat regions; see also **Table S3**). Therefore, we did not include any heterozygous SNPs or INDELs in our analyses, however, we have included a VCF file with these variants in the **Supplementary Materials**.

### 2.1 Genetic evidence for origin of C57BL/6 and C57BL/10 substrain differences

**Figure 1B** shows the relationships among C57BL/6 and C57BL/10 substrains based on historical records^37-39^. **Figure 1C** shows a dendrogram that was produced using SNPs, STRs and bi-allelic SVs. The number of concordant SNPs separating the substrains in the subtree under each branch from all the other substrains is indicated beside each branch. 342,002 SNPs (99.6% of non-monomorphic SNPs) agree with the dendrogram. Among 1,411 discordant SNPs no particular substrain pattern is dominant. Comparison of **Figure 1B** and **Figure 1C** shows that the records of the relationships among C57BL/6 and C57BL/10 substrains are consistent with our results.

### 2.2 Distribution of genomic variants across the genome

The distribution of variants across the genome is shown in **Figure 1D**. Several dense clusters of variants common in all categories (SNPs, INDELs, STRs and SVs) are evident (e.g. on chromosomes 4, 8, 11 and 13 for example). The non-uniformity of these polymorphisms is inconsistent with our expectation that polymorphisms among these substrains are due to new mutations and genetic drift. To further explore this observation, we examined the distribution of SNPs for each of the 14 different substrains; **Figure 1E** demonstrates that these clusters consist of a series of highly divergent haplotypes that differentiated the C57BL/6 and C57BL/10 lineages. In total 312,981 SNPs (89% of SNP variants detected in this study and 99.6% of the C57BL/6 vs C57BL/10 SNPs) reside in C57BL/10-specific clusters that represent just 5% of the genome (28 segments on 11 chromosomes: 1, 2, 4, 6, 8, 9, 11, 13, 14, 15, 18) with a SNP density of ∼1/425 bp. Across the remaining 95% of the genome SNPs do not appear to be clustered and have a density of ∼1/67,000 bp (more than 100-fold less dense). We found that many of the SNPs in these intervals are present in the Mouse Genome Informatics (MGI) database^40^, which further suggests that they are not due to new mutations in the C57BL/10 lineage. We used the MGI database to identify strains that were similar to these 28 segments. No single strain matched all 28 segments. However, **Figure S2** shows that for 24 of the 28 segments, at least one strain in the database has greater than 90% concordance (we only considered strains for which a minimum of 300 SNPs were available in that segment). Based on these data, we hypothesize that one or more inbred or outbred mice were accidentally or deliberately introduced into either the C57BL/6 or C57BL/10 lineage. Another possibility is that their last common ancestor was not fully inbred, and that these regions were differentially fixed after their separation. Most of the concordant strains that we identified in the MGI database have *domesticus* origin; however, two large segments on chromosomes 4 and 11 showed apparent *musculus* origin. We checked the subspecific origin of these 24 regions in C57BL/10J, C57BL/10ScNJ and C57BL/10ScSnJ reported in Yang *et. al*.^41^. The two large segments on chromosome 4 and 11 (with similarities to CZECHII/EiJ and MSM/Ms with *musculus* origins in MGI database) showed *musculus* origins as well. Among the remaining 22 segments all but three showed *domesticus* origins, and three segments (two on chromosome 4 and one on chromosome 11) showed some evidence of *musculus* origin.

Additionally, we identified 9,218 SNPs (2.6% of all SNPs) in which none of the C57BL/6 or C57BL/10 substrains matched the reference genome (mm10). One explanation is that some of these SNPs represent errors in mm10, perhaps related to the use of BACs or other technical issues^42^. Another explanation is that some of these SNPs represent true differences between the individuals used to generate mm10 and the individual C57BL/6J used for WGS in our study; we expect there should be some unfixed polymorphisms within C57BL/6J that exist at intermediate frequencies, meaning they will be observed in some individuals (e.g. the C57BL/6J individuals used to generate mm10) but not in other individuals (e.g. the C57BL/6J individual sequenced in our study)^43^. The distributions of other variants (INDELs, STRs and SVs) mirrored SNPs and are plotted in **Figure S3**.

### 2.3 Identification of candidate genomic variants causing differential gene expression

We found that 2,826 of 16,400 expressed genes (17.2%) were differentially expressed among the 14 substrains (FDR 0.05). We refer to these as DEGenes. We assumed that many of the DEGenes were due to local (*cis*) polymorphisms.

In order to identify genomic variants that might be causally related differential gene expression, we tested all identified variants (SNPs, INDELs, STRs and SVs) in the *cis-* window (1Mb upstream of gene start and 1Mb downstream of gene end) for association with the corresponding DEGene. Specifically, we tested the association between the *cis-* variants and the median of DEGene expression by a linear regression test, using Limix^44^. The resulting p-values are reported in the **Supplementary Material**.

As expected, all variants with the same strain distribution pattern have identical p-values in the association tests. For example, for the gene *Kcnc2*, which had significantly reduced expression in C57BL/6JEiJ (**Figure 2A)**, there was an equally strong correlation with four SNPs, one INDEL and one STR in the *cis*-region (**Figure 2B**), and many more variants outside the *cis*-region (**Figure S4**). The INDEL was annotated as a frameshift loss-of-function variant by Variant Effect Predictor (VEP)^45^, therefore, it had a strong prior to be the causal variant. Even in this small cohort of just 14 strains, we found a number of examples in which a variant within the *cis*-window had a strong prior and therefore appeared likely to explain a DEGene. We describe several such examples in the next section. However, for the majority of DEGenes, there were no polymorphisms that had strong priors, meaning that any of the variants with the smallest p-values in the *cis*-window, or a combination of them, or *trans*-acting variants elsewhere in the genome, could be causal.

**Figure 2.**
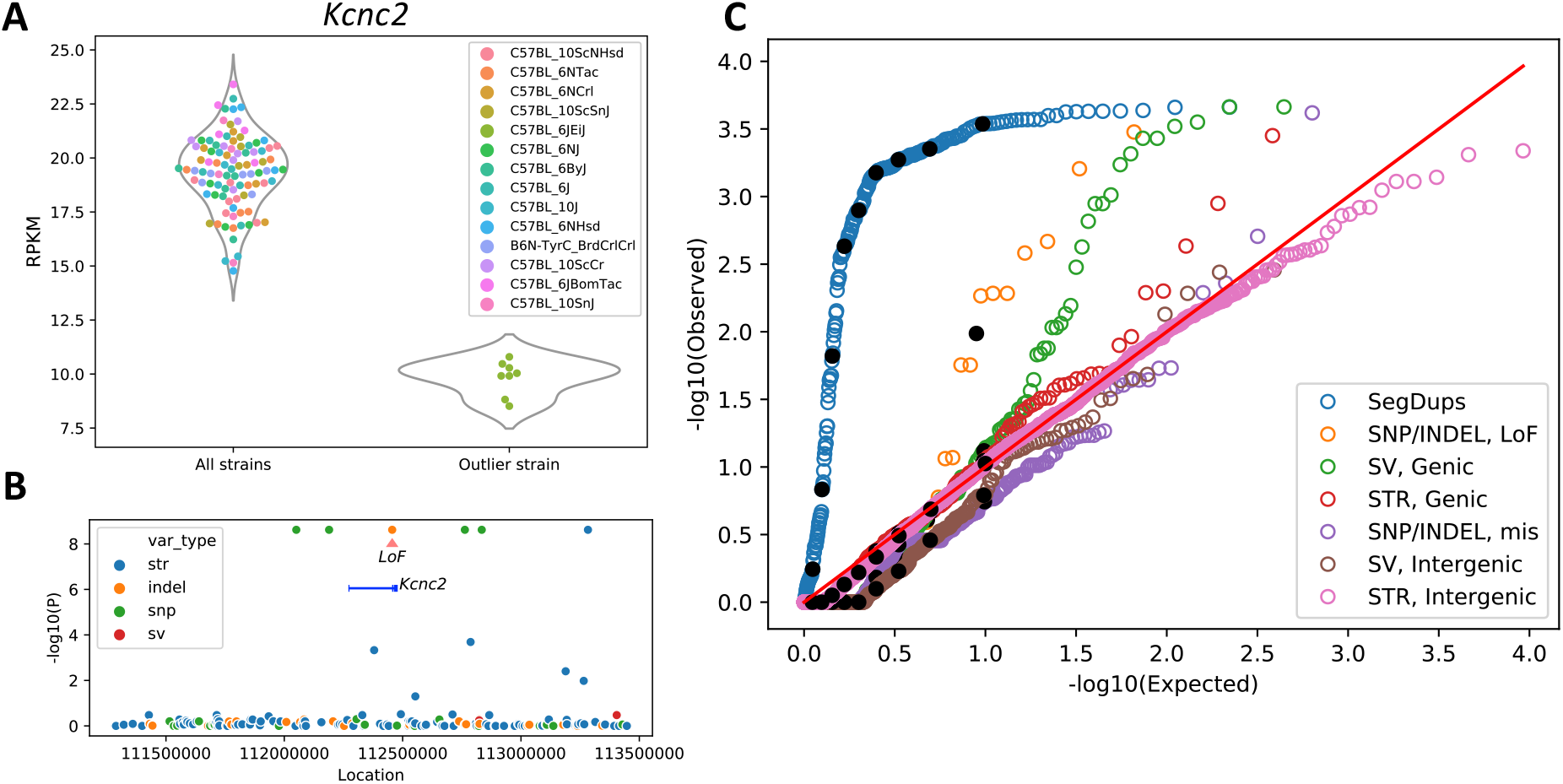
Association of gene expressions with genomic variants. **A:** Expression of *Kcnc2* is lower for C57BL/6JEiJ compared to the other substrains. **B:** *Cis*-variants of *Kcnc2* are tested for association with the median expression by the linear regression model. One INDEL is a frameshift loss-of-function variant and therefore has a strong prior to be the causal variant. In addition to that, four SNPs and one STR also have the same strain distribution pattern, and therefore the same -log10(p) value. For most of the DEGenes, no variant belonged to a class that had a strong prior for causality. In those cases, any of the variants with the smallest p-values (or a combination thereof, or more distant variants) might be causal (see also **Figure S4**). **C:** The distribution of the p-values of variants in different categories is compared against the uniform distribution in a QQ plot. A Linear Mixed Model is used with the Genomic Relatedness Matrix (GRM) as a random effect to control for population structure and the parental strain (C57BL/6 or C57BL/10) is used as a fixed effect to identify associations within C57BL/6 and C57BL/10 substrains. The black dots show the deciles of the data in each category. The SegDup category includes associations between the copy number variation of the DEGenes intersecting with SegDup regions (obtained by read depth across the segmental duplication regions of the reference genome) and the gene expression. Loss-of-function and missense mutations are two categories of SNP/INDELs. Genic SVs include those intersecting with gene features such as exons, TSS, UTRs, promoters, enhancers and introns, and genic STRs include those intersecting with exons, TSS, 5’UTRs and promoters. Intergenic SVs and STRs are those not intersecting with any gene features, and are paired with a gene with the closest TSS (see also **Table S2**).

### 2.4 Differential expression of genes is associated with multiple categories of functional variants

Genomic variants that disrupt protein coding exons or nearby *cis*-regulatory elements have the potential to cause differential gene expression. We investigated the causal role of variants in the *cis*-window by quantifying the strength of effects for multiple functional categories of variants. SNPs and INDELs were annotated using VEP^45^, which identified 183 (58 SNPs and 125 INDELs) loss-of-function variants (frameshift, stopgain or splice variant). We validated a random subset of these variants (13 SNPs and 29 INDELs) by Sanger sequencing (see STAR Methods and **Table S4**). All of the SNPs and INDELs showed 100% validation rates except for singleton INDELs, which showed much lower validation rate of 42%. SVs were annotated by intersecting with the gene features including exons, Transcription Start Site (TSS), Untranslated Regions (UTRs), promoters, enhancers, and introns. When an SV intersected with multiple types of functional elements, it was categorized according to the order mentioned above. The same gene annotations were applied to STRs. Intergenic SVs and STRs, which are defined as those that did not intersect with any gene features, were paired with the gene that had the nearest TSS. In addition, we assessed multiallelic copy number variation of genes by quantifying sequence coverage of all Segmental Duplications mapped to the mm10 reference genome^46^ that intersected with genes.

Genomic variants of the above categories that intersected with DEGenes were tested for association with gene expression by a Linear Mixed Model (LMM) using Limix^44^. We controlled for the complex relationships among inbred strains (a form of population structure) by using a Genomic Relatedness Matrix (GRM), derived from SNP genotypes, as a random effect, and parent strain (C57BL/6 or C57BL/10) as a fixed effect. **Figure 2C** shows the QQ plot for the p-values obtained from the data versus the uniform distribution. The black dots show the deciles of the data in each category. SegDups that intersected genes were strongly correlated with the expression of those genes, as would be expected for gene copy number variation. Loss-of-function SNPs and INDELs also showed a significant inflation of correlated DEGenes, followed by the genic SVs. The genic STRs showed a slight inflation, which was not as significant as other variant types. The missense SNPs, intergenic SV and intergenic STR p-values followed the uniform distributions.

For each category, the p-values obtained by the LMM model are corrected by the Benjamini-Hochberg procedure to obtain an FDR. We identified 53 significant (FDR < 0.05) associations between DEGenes and features, which are reported in **Table S2**. The majority of these associations (41 of 53 genes) reflected segmental duplications. In **Table S2** we report the genotype pattern in the substrains for each variant as well; notably, there are several clusters of significant associations with the same genotype pattern. For example, one extensive region on Chromosome 2 that clearly distinguishes C57BL/6NJ from all other substrains accounts for 18 of the 53 genes identified. Another cluster with a more complex pattern on Chromosome 4 accounts for 11 of the 53 identified genes.

### 2.5 Distinct mechanisms of differential gene expression caused by SVs

SVs can affect gene function by (1) varying the dosage of a full-length gene (2) deletion or insertion of exons producing alternative isoforms of a gene, or (3) rearrangement of the cis-regulatory elements of genes. For example, there are three copies of the gene *Srp54* in the mouse reference genome, but we found significant variability in the number of copies across the substrains; the number of copies was strongly associated with expression of *Srp54* (**Figure 3A**). Thus, in this example, copy number variation in segmental duplication regions is the likely cause of differential gene expression. An example of a SV that likely impacts expression is the *Lpp* gene. The *Lpp* gene has a tandem duplication of the first two exons in two substrains (C57BL/10ScCr and C57BL/10ScNHsd) that creates two copies of the TSS, which probably accounts for its ∼2-fold increased expression (**Figure 3B**).

**Figure 3.**
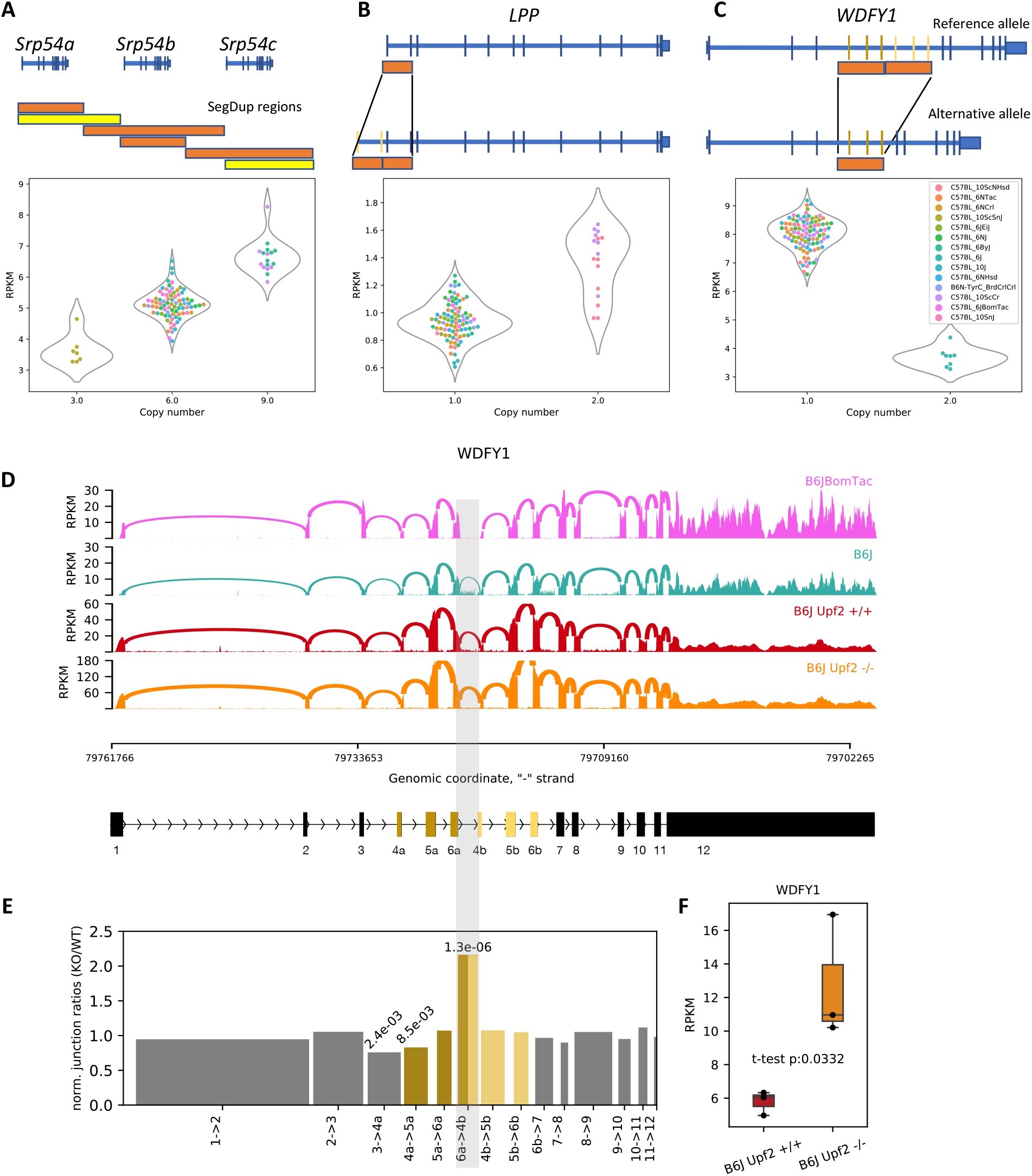
Structural variations affecting expression of *Srp54, Lpp* and *Wdfy1*. **A**: Variation in copy number in a SegDup region is associated with expression of *Srp54*. The read coverage in these SegDup regions is used to infer the number of copies of the intersecting genes. The bars with similar lengths and colors indicate corresponding SegDup regions. Yellow bars indicate more than 98% sequence similarity, and orange bars indicate more than 99% sequence similarity. **B**: A duplication involving the first two exons of *Lpp* in two substrains of C57BL/10 is associated with an increase in *Lpp* expression. Duplication of the TSS and the promoter site is the most likely cause. **C**: A segmental duplication region which intersects with three exons in *Wdfy1*, and is present in C57BL/6J, is associated with reduction of expression of *Wdfy1* in C57BL/6J. All the other substrains lack this duplication. **D**: Sashimi plots^51^ for C57BL/6BomTac (a closely related substrain to C57BL/6J), C57BL/6J, *Upf2*^+/+^, and *Upf2*^-/-^ cell lines from C57BL/6J highlighting the junction between 6a and 4b exons across the segmental duplication region. Since C57BL/6BomTac lacks the duplication, it does not have any junctions between those exons, while the relative number of junctions in the *Upf2*^-/-^ cell line is significantly larger than the other wild type C57BL/6J samples. **E**: The bar plot shows the ratio of the normalized number of junctions between 6a and 4b exons (normalized by the total number of junctions in each sample) in *Upf2*^-/-^ over *Upf2*^+/+^ cell lines. It shows a significant increase in the relative number of junctions between the two segmental duplications in the *Upf2*^-/-^ cell line. The numbers on top of the bars show p-values obtained by the Chi-squared test. **F**: The expression level of *Wdfy1* in the *Upf2*^-/-^ cell line is significantly higher than the *Upf2*^+/+^ cell line. This supports our hypothesis that the reduction of gene expression in C57BL/6J is due to the nonsense mediated decay (NMD) mechanism.

An intriguing example of altered expression caused by a SV is the *Wdfy1* gene. This gene has a tandem duplication of exons 4-6 in the C57BL/6J substrain, which is also present in the mm10 reference genome. We found this duplication is associated with a paradoxical decrease in *Wdfy1* gene expression (**Figure 3C**), a result that could potentially be explained by nonsense-mediated RNA decay (NMD) of transcripts that contain duplications of exons 4-6. NMD is a highly conserved pathway that promotes the turnover of mRNAs harboring premature termination codons, including those generated by frameshifts^47^. We found no evidence of unproductive transcripts *of Wdfy1* in Gencode VM23. However, we reasoned that if a major spliced isoform of *Wdfy1* contains tandemly duplicated exons, it could be detected in cells that are deficient in NMD. RNA-Seq analysis of NMD-deficient (*Upf2*^-/-^) C57BL/6J ES cells showed that *Wdfy1* expresion was increased ∼2-fold compared to control (sibling) C57BL/6J ES cells (**Figure 3F**). Analysis of splice junctions from RNA-Seq confirmed the existence of an aberrant isoform in all C57BL/6J lines that includes splicing of exon 6 to the downstream (duplicated) exon 4 (**Figure 3D**), which we refer to as the “ 6a->4b” junction. This splice junction was unique to C57BL/6J strains, and the ratio of 6a->4b to all splice junctions was increased by ∼2-fold in the *Upf2*^-/-^ ES cells relative to control ES cells (**Figure 3E**). These results demonstrate that certain isoforms transcribed from *Wdfy1* in C57BL/6J mice are degraded by NMD, and the transcripts that are retained are alternative splice forms that exclude the 6a->4b junction.

## 3. Discussion

We have performed a large-scale multi-omics analysis of 14 C57BL substrains. We identified 352,631 SNPs, 109,096 INDELs, 150,344 STRs and 3,425 SVs;furthermore, of the 16,400 genes that were expressed in the hippocampus, 2,826 were significantly differentially expressed (FDR<0.05). Unexpectedly, many of the polymorphisms that differentiated the C57BL/6 and C57BL/10 substrains were concentrated in a few haplotypes, comprising just 5% of the genome. These polymorphisms appear to be due to either introgression of an unrelated individual, or incomplete inbreeding at the time that the C57BL/6 and C57BL/10 lineages diverged. Setting these introgressed regions aside, we tried to identify variants that were causally related to differential gene expression by focusing on the *cis* regions around DEGenes. This allowed us to identify 53 genes in which a variant with high prior probability to be causal was significantly associated with gene expression. While the majority of these 53 instances were caused by segmental duplications, several of which spanned many adjacent genes, a smaller proportion were due to SVs and INDELs (see **Table S2**). Inflation of test statistics for these categories of variants further underscores their likely causal roles; a relaxed FDR threshold would have identified more than 53 variant/DEGene associations.

An unexpectedly large subset of variants (89% of all SNPs) were concentrated in 28 highly diverged haplotypes that were present in all C57BL/10 strains and represented just 5% of the genome. These dense clusters of genetic variation (1 SNP/425 bp) perfectly differentiated C57BL/10 from C57BL/6, and likely reflect introgression from another strain. Intriguingly, the smaller haplotypes appeared to be of *domesticus* origin, and were similar to haplotypes found in multiple non-C57BL inbred strains. The two largest haplotypes appeared to be of *musculus* origin and were also similar to multiple non-C57BL inbred strains. The exact sequence of events that led to this situation are impossible to deduce, but these patterns are clearly due to breeding events (intentional or accidental) rather than spontaneous mutations; this conclusion is based on several observations: 1) the density of the polymorphisms, 2) the abrupt boundaries of the regions/haplotypes and 3) the fact that the SNPs in these introgressed regions are found in other inbred strains, which would not be the case if they were due to spontaneous mutations. A previous microarray study performed on 198 inbred mouse strains also identified SNP differences between C57BL/6J and three C57BL/10 substrain (C57BL/10J, C57BL/10ScNJ and C57BL/10ScSnJ) for all the 28 introgressed segments that we identified^41,48^; however, that study did not highlight the significance of that finding, and did not have sufficiently dense coverage to define the boundaries of the introgressed regions. While a majority of C57BL/10-specific genetic variants lie within these introgressed regions, they contained only a small fraction (∼13%) of DEGenes; however, given that the introgressed regions represent only 5% of the genome, this is still more than a 2-fold greater density of DEGenes that would be expected if they were randomly distributed across the genome.

Outside of these apparently introgressed regions we identified 37,745 SNPs that were distributed throughout the genome in a Poisson fashion with more than 100-fold lower density (∼1 SNP/67,000 bp). These SNPs are apparently due to the accumulation of new mutations and their identification was the original goal of our study. Dendrograms based on these SNPs recapitulated the historically recorded relationships among the substrains (**Figure 1B**). For the relatively large number of DEGenes (>2,000) that were located outside of the introgressed regions, we considered the association between different categories of nearby (*cis*) variants, and expression of DEGenes. Variable copy number segmental duplicated regions were shown to be highly enriched for significant associations as were genic SVs and loss of function SNP/INDELs (**Figure 2C**).

We presented several examples to highlight how different classes of variants underlie DEGenes. For example, variable copy number segmental duplications led to both increased and decreased expression of *Srp54* (**Fig 3A**). In another example, duplication of transcription start sites led to increased expression of *Lpp* (**Fig 3B**). In the case of *Wdfy1*, duplication of several exons led to down-regulation of expression (**Fig 3C**), which we showed was due to NMD-mediated mRNA decay (**Fig 3D**). *Wdfy1* was previously reported to be differentially expressed between C57BL/6J and C57BL/6NCrl, and was identified as one of the candidate genes for reduced alcohol preference in C57BL/6NCrl^49^. This gene is also within the QTL named *Emo4* (location: Chr1:68,032,186-86,307,305 bp)^50^; mice which are homozygous for C57BL/6J allele are more active in the open field test. Whether *Wdfy1* is actually the cause of either association cannot be resolved by our study.

Despite the numerous examples in which likely causal variants were identified, a majority of the causal variants underlying DEGenes remain unknown. Many are likely to be due to variants in regulatory regions that have not been distinguished from other nearby variants with the same strain distribution pattern (and thus identical p-values). Although we focused on the possibility that DEGenes were due to nearby variants (*cis*-eQTLs), the large fraction of differentially expressed genes (17.2% of all expressed genes) could indicate that many DEGenes are due to *trans*-eQTLs. Producing crosses between pairs of strains will be necessary to address the relative importance of *cis-* versus *trans*-eQTLs in the observed DEGenes; it is possible that such crosses could identify one or more major *trans*-regulatory hot spots.

Our results create a resource for future efforts to identify genes and causal polymorphisms that give rise to phenotypic differences among C57BL strains using the increasingly popular reduced complexity cross (**RCC**) approach in which two phenotypically divergent nearly isogenic inbred substrains are crossed to produce an F2 population^24^. Because of the low density of polymorphisms, identifying the causal allele is much more tractable. For example, the gene *Cyfp2* was identified as the cause of differential sensitivity to cocaine and methamphetamine in a cross between C57BL/6J and C57BL/6N^21^. In the **Supplementary material** we have provided genomic variants (SNPs, INDELs, STRs and SVs), differentially expressed genes in the hippocampus, as well as association tests between DEGenes and nearby variants. In addition, we have provided the VEP annotated SNP/INDELs, which distinguish loss of function, missense and synonymous mutations. Our data also identify some regions that have a high density of polymorphisms that may complicate the RCC approach. For example, phenotypic differences between C57BL/6 and C57BL/10 strains might frequently map to the introgressed regions, which have a high density of polymorphisms that would significantly hinder gene identification and negate many of the advantage of RCCs. Furthermore, crosses between two C57BL/6 or between two C57BL/10 strains may map to large segmental duplication regions such as those on Chromosomes 2 and 4 (see **Figure 1E** and **Table S2**), which would again hinder gene identification. Thus, one key observation from this study is that genetic differences among and between C57BL/6 and C57BL/10 strains are not uniformly distributed. Furthermore, our study used a single individual to represent each strain for whole genome sequencing. Therefore, we did not explore the extent to which the polymorphic regions we identified may be segregating versus fixed within each inbred strain. If some of these polymorphic regions are not fixed, it would further complicate the analysis of RCCs.

Whereas the RCCs represent a forward genetic approach (starting with a phenotypic difference, searching for the genetic cause), another novel application of our dataset would be to select two strains that are divergent for a coding or expression difference and to use that cross to study gene function. This reverse genetic approach (starting with a genetic difference, searching for the phenotypic consequences) has not been attempted using closely related substrains, but is conceptually similar to characterization of a knockout mouse. This approach is limited by the available polymorphisms. Although it would be necessary to account for the impact of linked polymorphisms, most of the polymorphisms would be unlinked and would not confound the interpretation of results.

In summary, we have created a dataset that elucidates the differences among C57BL strains and can be used for both forward genetic (RCC) and reverse genetic approaches. We identify previously unknown introgressed segments that differentiate the C57BL/6 and C57BL/10 lineages. Our results can also be used to explore mutational processes and highlight the tendency of inbred strains to change over time due to the accumulation of new mutations and genetic drift.

## Supporting information

Key resources table

## Acknowledgements

M.M., Y.R., C.L.S.P., A.W. and A.P. were supported by P50DA037844. J.S. was supported by MH113715. Y.R. was supported by T32MH018399 and A.W. was supported by T32MH020065.

## Author Contributions

A.P. designed the study. C.L.S.P. performed the animal breeding, dissection and the preparation of WGS and RNA-Seq libraries. Y.R. and A.W. performed initial analyses of the WGS and RNA-Seq data. M.M. carried out all statistical genetic and functional genomic analyses. M.G. and S.S. performed STR calling. M.M. and J.S. performed SV calling and analysis of SV eQTLs. M.W. developed the *Upf2* mouse model, A.S. derived the ES cells and performed the corresponding RNA-Seq. M.M., J.S., A.P. wrote the paper.

## STAR Methods

### RESOURSE AVAILABILITY

#### Lead contact

Further information and requests for resources should be directed to and will be fulfilled by the Lead Contact, Milad Mortazavi.

#### Material availability

This study does not generate new unique reagents.

#### Data and code availability

- RNA-Seq and WGS short read raw data have been deposited at NCBI SRA and are publicly available as of the date of publication. Accession numbers are listed in the key resources table. All processed data have been deposited at Mendeley and are publicly available as of the date of publication. DOIs are listed in the key resources table.
- All original code has been deposited at Zenodo and is publicly available as of the date of publication. Accession numbers are listed in the key resources table.
- Any additional information required to reanalyze the data reported in this paper is available from the lead contact upon request.

### EXPERIMENTAL MODEL AND SUBJECT DETAILS

#### Mice

We obtained a panel of 14 C57BL substrains from four vendors. The panel included 9 C57BL/6 substrains: C57BL/6J, C57BL/6NJ, C57BL/6ByJ, C57BL/6Ntac, C57BL/6JbomTac, B6N-TyrC/BrdCrCrl, C57BL/6NCrl, C57BL/6NHsd, C57BL/6JeiJ, and 5 C57BL/10 substrans: C57BL/10J, C57BL/10ScCr, C57BL/10ScSnJ, C57BL/10SnJ, C57BL/10ScNHsd (**Table 1**). All of the substrains were bred for one generation at the University of Chicago before tissue was collected for whole genome sequencing and RNA-sequencing; this avoided gene expression differences that were secondary to environmental differences among the four vendors. Mice were ordered in November 2014, arrived at University of Chicago in December 2014, and started breeding in January 2015. Tissues were extracted from the first generation at the age of 50-60 days old for RNA-sequencing. All procedures were approved by the University of Chicago IACUC. One hundred and ten male mice in total, with six to eleven mice per substrain, were chosen for RNA-sequencing from hippocampus, and one male mouse per substrain was chosen for whole genome sequencing from spleen (**Figure 1A**).

#### Whole-genome sequencing (WGS)

DNA from one male animal per substrain (n=14) was extracted from spleens using a standard “ salting-out” protocol. Sequencing libraries were prepared using a TruSeq DNA LT kit, as per the manufacturer’s instructions. Subsequently, sequencing data was generated by Novogene at an average depth of ∼30X coverage on an Illumina HiSeq X Ten (paired-end 150bp) (**Table 1**).

#### RNA-sequencing

Total RNA was extracted from 110 hippocampal samples using Trizol reagent (Invitrogen, Carlsbad, CA). RNA was treated with Dnase (Invitrogen) and purified using Rneasy columns (Qiagen, Hilden, Germany). RNA-sequencing library prep and sequencing was performed by the University of California San Diego Sequencing Core using Illumina TruSeq prep and Illumina HiSeq 4000 machine (single-end 50bp; **Table 1**).

#### Nonsense mediated decay assay

To determine whether SVs of the *Wdfy1* gene in C57BL/6J create novel mRNA isoforms that are degraded by the Nonsense-Mediated Decay (NMD) pathway, we performed RNA-Seq on mouse embryonic stem cells (mESCs) from a *Upf2*^-/-^ strain of C57BL/6J that has impaired NMD and control mouse mESCs from C57BL/6J. Samples with an RNA integrity index of >8 (as determined by a BioAnalyzer) were used for RNA-Seq analysis. The University of California San Diego Sequencing Core performed library preparation using ribosomal RNA depletion protocol followed by paired-end sequencing (100 cycles) using a HiSeq4000.

### METHOD DETAILS

#### RNA data processing

Reads were mapped to mouse reference transcriptome (mm10) using the splice-aware alignment software HiSat2^53^, and counts were normalized using HTSeq^54^. Only genes that had at least one Count Per Million (CPM), for at least two samples were included in our analysis. We further removed four outlier samples identified by PCA analysis. This left us with gene expression data for 16,400 genes across 106 samples in 14 substrains. To identify Differentially Expressed Genes (DEGenes) we performed analysis of variance using the *anova* function in R, and adjusted the p-values by computing the false-discovery rate (FDR) using the *p*.*adjust* function in R, with the Benjamini-Hochberg procedure. We obtained 2,826 DEGenes among C57BL/6 and C57BL/10 substrains combined, 1,210 DEGenes within C57BL/6, and 104 DEGenes within C57BL/10 substrains with FDR<0.05.

Reads from three replicates of *Upf2*^-/-^ samples and three controls from the NMD assay were mapped to the mouse reference genome (mm10) by HiSat2^53^, and counts were normalized using HTSeq^54^. We kept all genes with CPM>1 and normalized the counts with *edgeR*^*55*^ function in *R*, however, we only analyzed *Wdfy1* expression in an effort to detect differences in NMD between *Upf2*^-/-^ and control samples.

#### SNPs and INDELs

We used SpeedSeq^56^ to process the WGS paired-end reads. SpeedSeq uses BWA-mem (v.0.7.8) to map the reads to the mm10 reference genome, SAMBLAST^57^ to mark duplicates, Sambamba^58^ to sort the BAM files, and FreeBayes^59^ to jointly call SNPs and INDELs. INDELs are defined as insertions or deletions which are relatively short in length. The length range for the detected INDELs in our study is between one and 64 base pairs, which is approximately the lower bound for SV length scales. We restricted our analysis to variants that were fixed within individual substrains by including homozygous SNPs and INDELs only, resulting in a callset consisting of 352,631 SNPs and 109,096 INDELs. To assess validation rates of these variants, we utilized the RNA-Seq data to validate WGS variants in protein coding regions of the genome. Reads from samples in each substrain were combined and GATK best practices pipeline for RNA-Seq variant calling^52^ was used. Average genome-wide read coverage of RNA-Seq data for fourteen substrains ranged from 1.81x to 3.71x with a median of 3.08x. The reads were mapped to the reference genome (mm10) by STAR with 2-pass option^60^. Subsequently, SplitHCigarReads command was used, followed by the base recalibration step. Afterwards, HaplotypeCaller was run on each substrain separately.

In each substrain, we validated a subset of WGS variants which were in the protein coding regions of the mouse genome (exons and UTRs obtained from Ensembl annotations), and had at least 3x coverage in our RNA-Seq data (24,260 SNPs, 5,637 INDELs).

Genotypes from the WGS data in the protein coding regions were compared with variants detected in RNA-Seq data, and validation rates for SNPs and INDELs as well as different categories of variants including C57BL/10-specific, singleton and monomorphic variants were shown in **Figure S1** and **Table S3**. Overall, we observed 97% validation rate for the SNPs and 67% validation rate for INDELs. Among INDEL categories, C57BL/10-specific and monomorphic INDELs were among highest validation rates (72%) and singletons showed the lowest validation rate (52%). Information about the position of the validated SNPs and INDELs can be found in the **Supplementary material**.

In order to validate the 183 (58 SNPs and 125 INDELs) predicted loss-of-function variants by VEP^42^, we performed Sanger sequencing for 13 randomly selected SNPs (7 C57BL/10-specific, 5 singleton, 1 monomorphic) and 29 randomly selected INDELs (9 C57BL/10-specific, 12 singleton, 8 monomorphic) (see **Table S4**). For each locus, we genotyped one sample from a randomly selected C57BL/10 substrain for C57BL/10-specific and monomorphic categories, or the substrain which had the singleton variant for the singleton category, and one C57BL/6J sample for control. The variants are provided in the **Supplementary material**.

When computing the identity-by-state (IBS) matrix for dendrograms, we LD-pruned the SNP panel with Plink^61^ (--indep-pairwise 50 5 0.5) yielding 16,739 SNPs. This pruned SNP set was augmented by STRs and all bi-allelic SVs, followed by computing the distance matrix with *dist* and plotting the dendrograms with *hclust* in *R* v3.6.1.

#### Short Tandem Repeat (STR)

We used HipSTR v0.6 with default parameters^62^ to call STRs from mapped reads using the mm10 reference STR set available from the HipSTR website^63^. The reference STR set was generated using Tandem Repeats Finder^64^ allowing a maximum repeat unit length of 6bp. STRs for the substrains were jointly genotyped on a single node of a local server in batches of 500 STRs. Resulting VCF files from each batch were merged to create a genome-wide callset in VCF format. We filtered out calls with missing genotypes, as well as calls with reference alleles for all substrains, resulting in a total of 150,344 polymorphic STRs. The STR calls are available in the **Supplementary material**.

#### Structural Variations (SV)

SVs were detected using a combination of approaches. First, we called SVs with LUMPY^65^ and CNVnator^66^, two complementary methods that rely on discordant and split read signals or coverage respectively. Second, because SV calling accuracy by the above methods is low in regions that are dense in segmental duplications, copy number variation within annotated segmental duplications was quantified directly from coverage, and these coverage values were used for the correlation of gene copy numbers with gene expression.

We filtered out SV calls that overlapped 50% or more with the gap regions of the mouse reference genome, as well as the calls with length smaller than 50 bp and larger than 1 Mbp. A more stringent >1000 bp length filter was applied to CNVnator calls. We then filtered out non-homozygous calls and calls that were homozygous for the alternative allele in all substrains.

Concordant calls from LUMPY and CNVnator with 50% or greater reciprocal overlap and the same genotypes were merged and the breakpoints reported by LUMPY were used. Consensus calls that overlapped with annotated segmental duplications (SegDup) in the reference genome were excluded, and instead SegDup copy number was assessed directly from read depth signal using mosdepth v0.2.6^67^ with window size 100 bp. SegDup annotations from the mm10 genome with at least 98% similarity were intersected with gene annotations, and the median read coverage across SegDups which intersect with genes was normalized by the median coverage of the corresponding chromosome. These normalized coverage values were used to correlate gene copy numbers with gene expression. The final set of SVs included 3,425 deletions, duplications and inversions in nine C57BL/6 and five C57BL/10 substrains. The distribution of SVs in each category and substrain is summarized in **Table S1**. The VCF file of the SV calls, and the read coverage data for the SegDup regions are provided in the **Supplementary material**.

### QUANTIFICATION AND STATISTICAL ANALYSIS

The details regarding association analyses are presented in the results sections as well as in the figure legends, and the software information is presented in the key resources table. P-value of 0.05 after correction for multiple testing (Benjamini-Hochberg procedure) was used to identify significant associations. The t-test and Chi-squared tests in Figure 3 was performed by scipy module in python v3.

## Supplemental information

**Figure S1.**
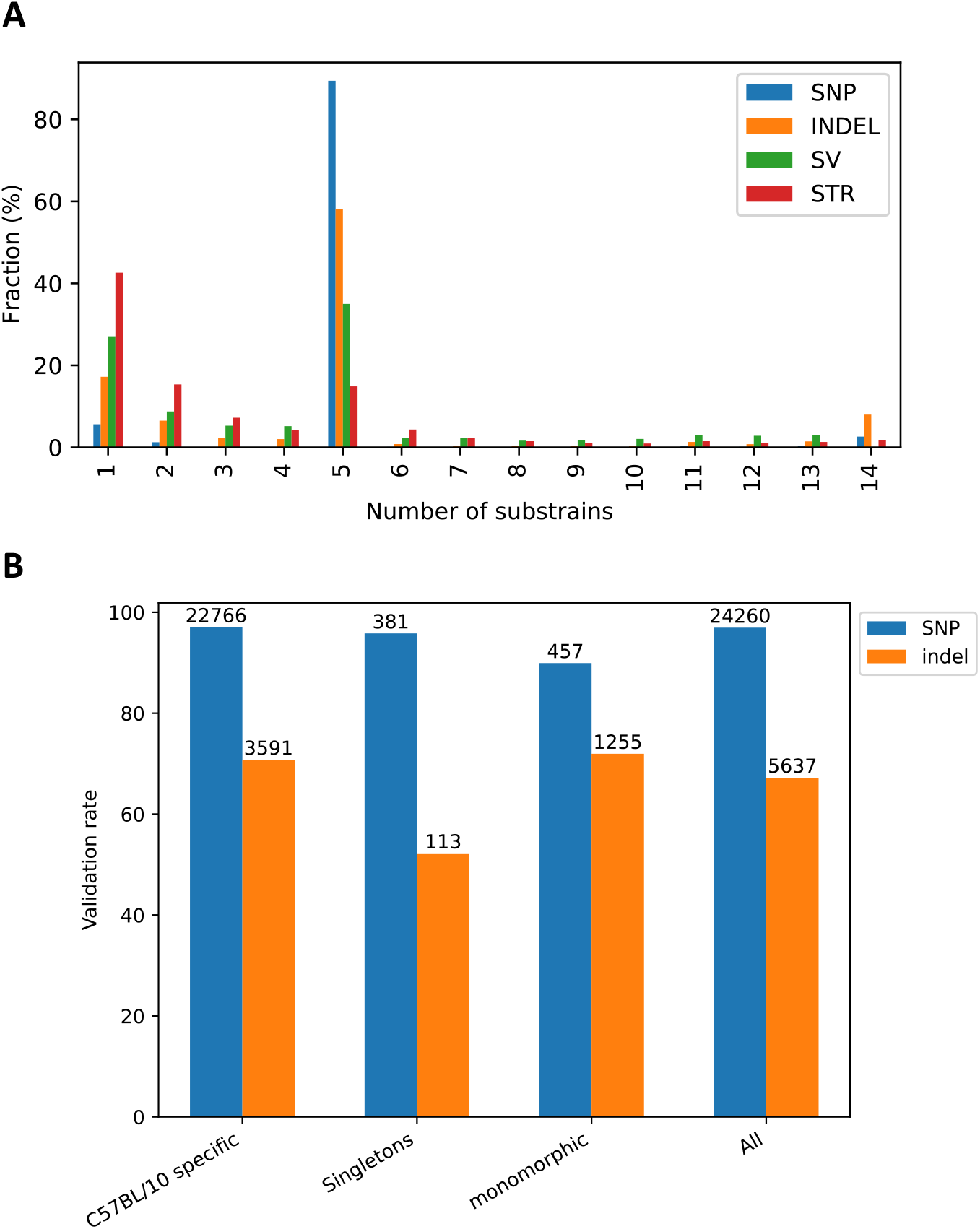
Variant frequency and validation rates, related to STAR Methods. **A:** Fraction of variants observed in substrains (five C57BL/10 and nine C57BL/6 substrains) in each variant category. The spike at 5 reflects polymorphisms that separated C57BL/10 (n=5) from C57BL/6 (n=9) substrains. The smaller spike at 14 represents instances where none of the substrains (including C57BL/6J, which is the basis for mm10) matched the mm10 reference genome. **B:** Validation rates of WGS variants in the protein coding regions using RNA-Seq data. WGS SNPs and INDELs which intersect with protein coding exon and UTR annotations (from Ensembl) and have at least 3X coverage in RNA-Seq dataset are considered for validation. Variants from RNA-Seq data were called by GATK best practices^52^ for each substrain separately (see also STAR Methods). Validation rate between different categories of variants are compared. The total number of variants in each category is indicated on top of each bar. Overall 97% of all SNPs and 67% of all INDELs were validated using RNA-Seq data.

**Figure S2.**
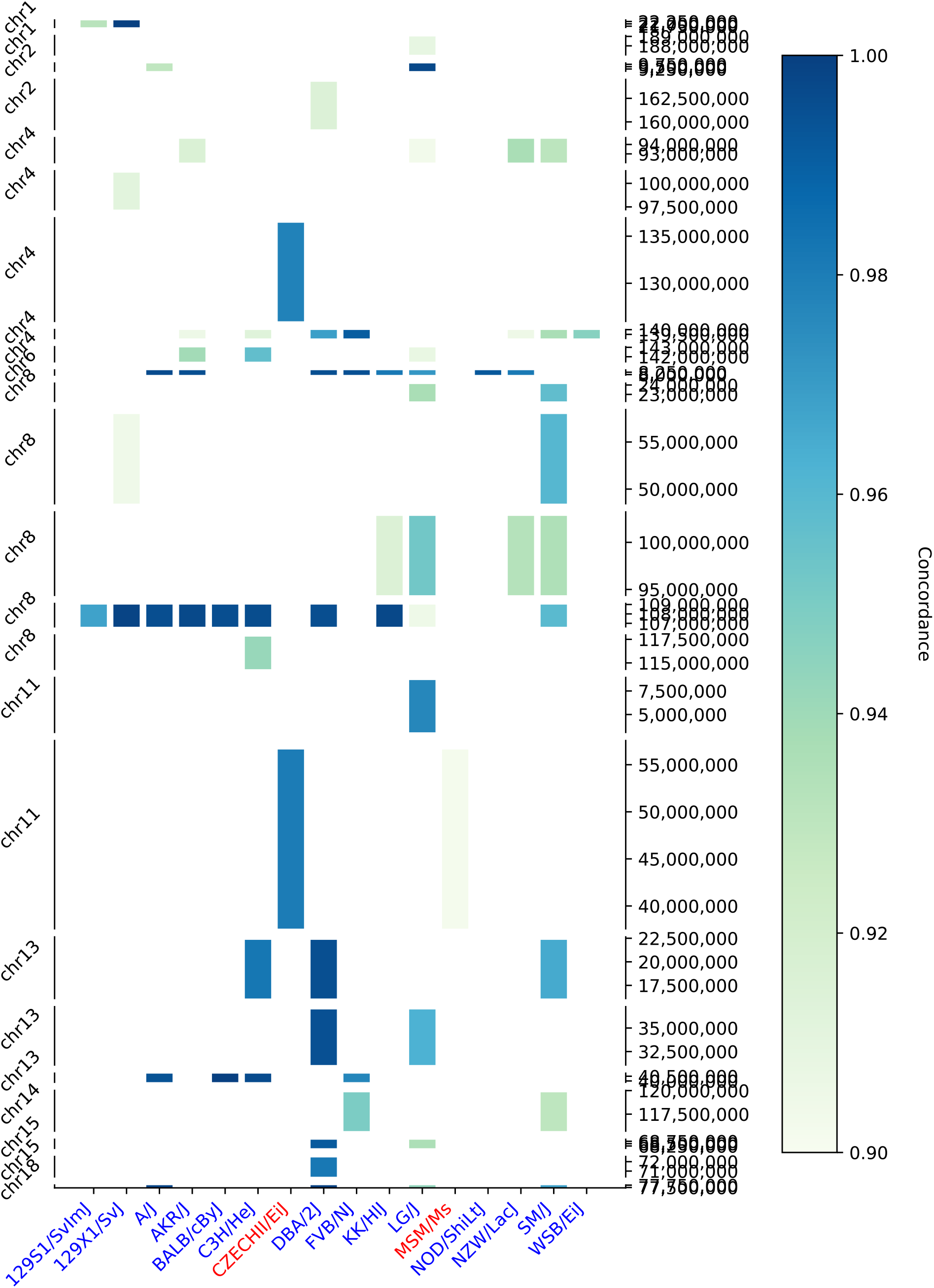
Concordance of C57BL/10-specific hotspot segments with other mouse strains, related to Figure 1. Concordance of 24 C57BL/10-specific haplotypes (SNP hotspots) with SNPs of other strains from *domesticus* and *musculus* origin. Y-axis shows segments with C57BL/10-specific SNP hotspots. X-axis shows strains that have at least 300 common loci and at least 90% concordance with C57BL/10-specific SNPs in each segment. The SNP data for the strains is obtained from MGI^40^. The segments are color coded with the concordance value. The strain labels on the x-axis are color coded with blue: *domesticus* origin, and red: *musculus* origin^41^.

**Figure S3.**
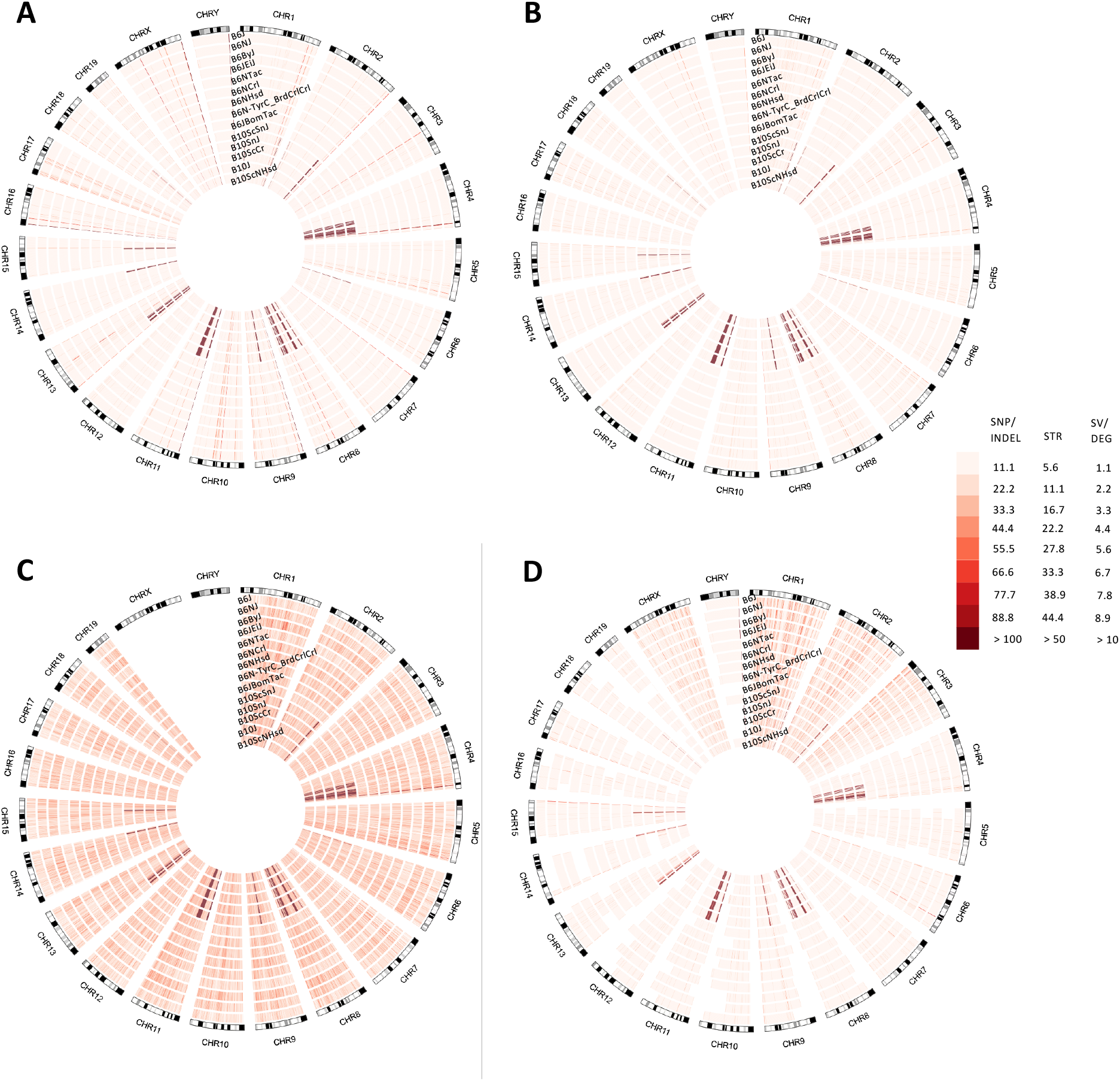
Distribution of genomic variations in 14 C57BL substrains, related to Figure 1. **A**: SNP distribution, **B**: INDEL distribution, **C**: STR distribution, and **D**: SV distribution for nine C57BL/6 and five C57BL/10 substrains show clusters of variants that are specific to C57BL/10 substrains on chromosomes two, four, eight, nine, eleven, thirteen, fourteen and fifteen.

**Figure S4.**
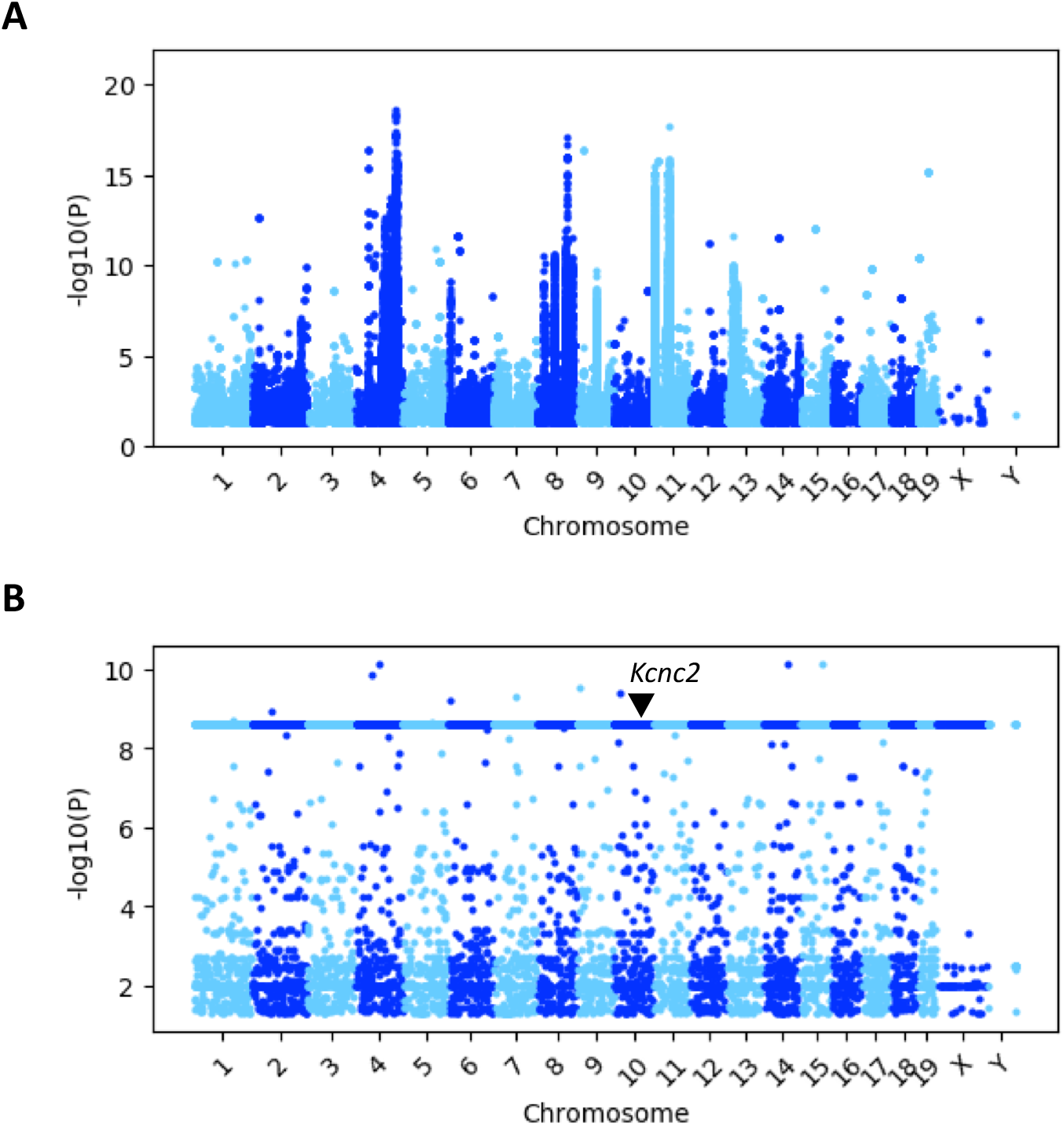
Association of all genomic variants and expression of DEGenes, related to Figure 2. Association tests of DEGene expressions of C57BL/6 and C57BL/10 substrains with all genomic variants (SNPs, INDELs, STRs and SVs) was performed by linear regression model with Limix^44^ **A**: Association of DEGene expressions with all variants (SNPs, INDELs, STRs and SVs) in the *cis*-region defined as 1Mb upstream and 1Mb downstream of the DEGene. The p-values are plotted at the genomic locations of the corresponding DEGenes. **B**: Association of *Kcnc2* expression with all genomic variants across the genome shows that variants with the same strain distribution pattern have identical p-values. The flat horizontal line at about -log10(p)=8.4 reflects features that have the same strain distribution pattern and therefore all yield identical p-values when tested for association with the gene expression data.

**Table S1.** Number of SVs found in C57BL substrains, related to Methods details.

|  |  | CNVnator |  |  | Lumpy |  |
| --- | --- | --- | --- | --- | --- | --- |
| Strain | Substrain | DEL | DUP | DEL | DUP | INV |
| C57BL/6 | C57BL/6J | 27 | 461 | 11 | 97 | 2 |
|  | C57BL/6NJ | 5 | 456 | 65 | 105 | 2 |
|  | C57BL/6ByJ | 11 | 426 | 82 | 90 | 5 |
|  | C57BL/6JeiJ | 14 | 469 | 42 | 91 | 4 |
|  | C57BL/6Ntac | 27 | 465 | 64 | 77 | 4 |
|  | C57BL/6NCrl | 9 | 420 | 66 | 92 | 2 |
|  | C57BL/6NHsd | 127 | 488 | 73 | 86 | 3 |
|  | B6N-TyrC/BrdCrCrI | 7 | 445 | 63 | 96 | 3 |
|  | C57BL/6JbomTac | 13 | 447 | 50 | 78 | 3 |
| C57BL/10 | C57BL/10ScSnJ | 95 | 430 | 1204 | 129 | 15 |
|  | C57BL/10SnJ | 89 | 420 | 1195 | 125 | 13 |
|  | C57BL/10ScCr | 269 | 481 | 1196 | 103 | 15 |
|  | C57BL/10J | 74 | 437 | 1193 | 115 | 16 |
|  | C57BL/10ScNHsd | 72 | 426 | 1192 | 123 | 14 |
|  | TOTAL | 448 | 1308 | 1369 | 279 | 21 |

**Table S2.**
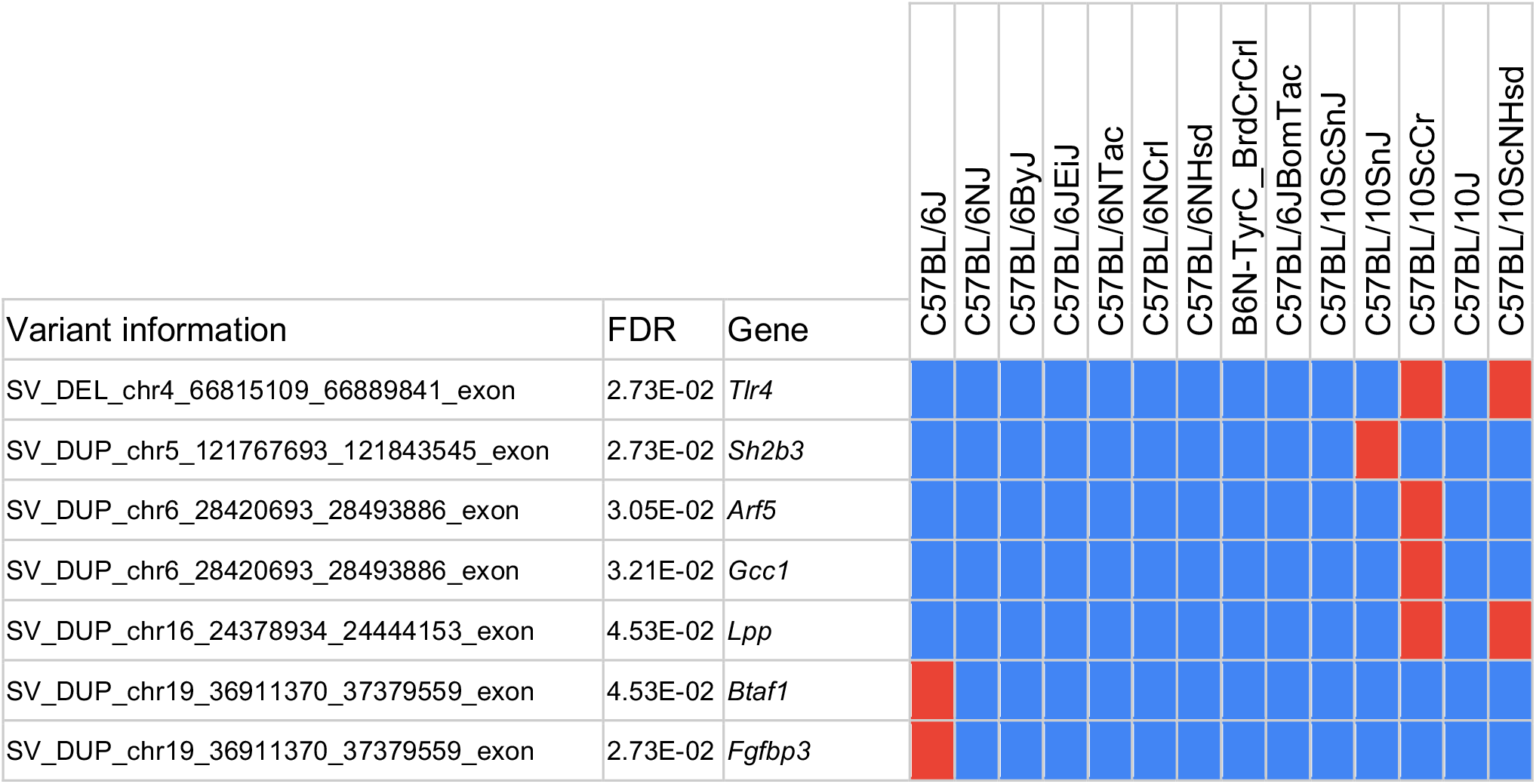
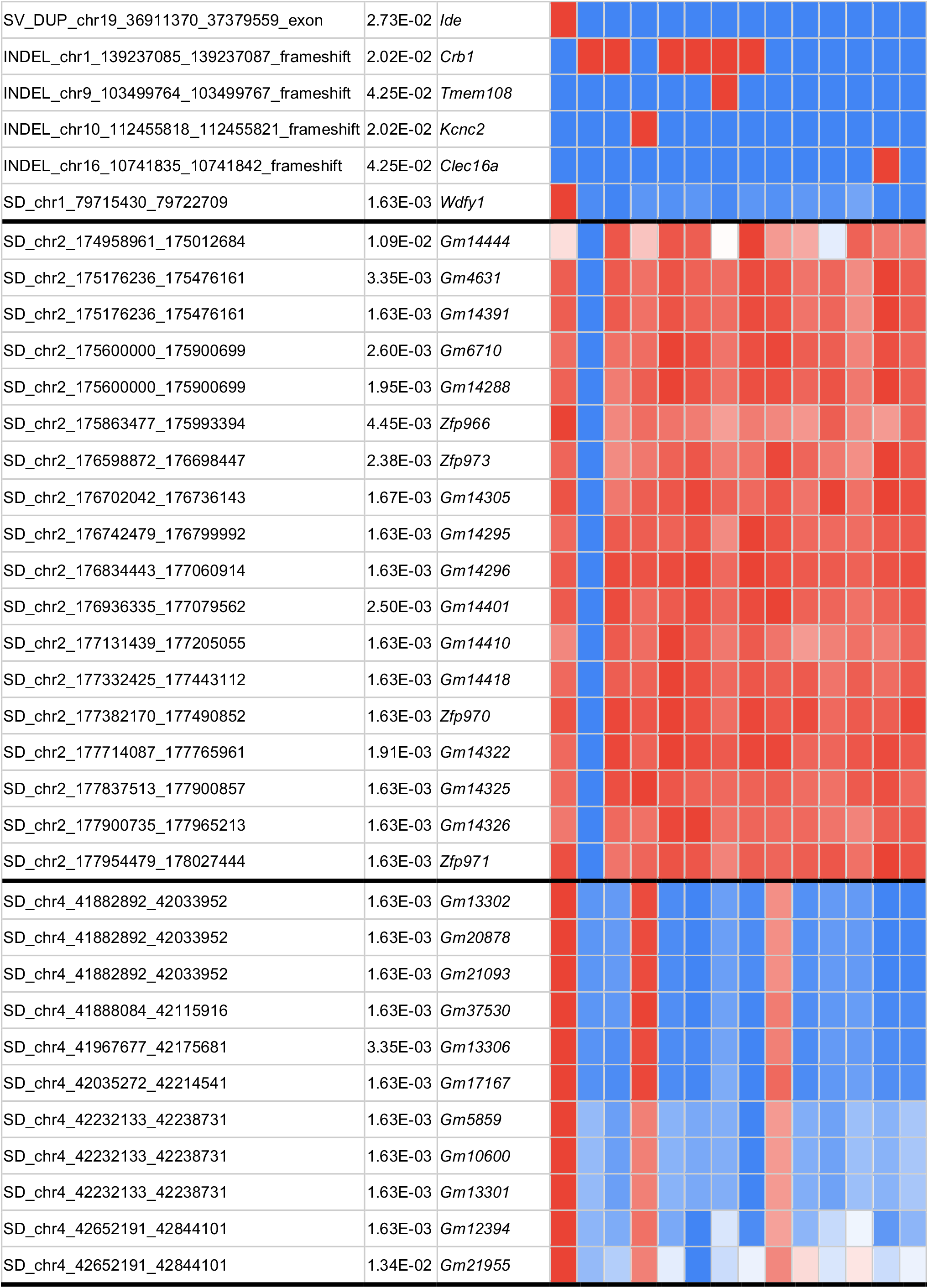
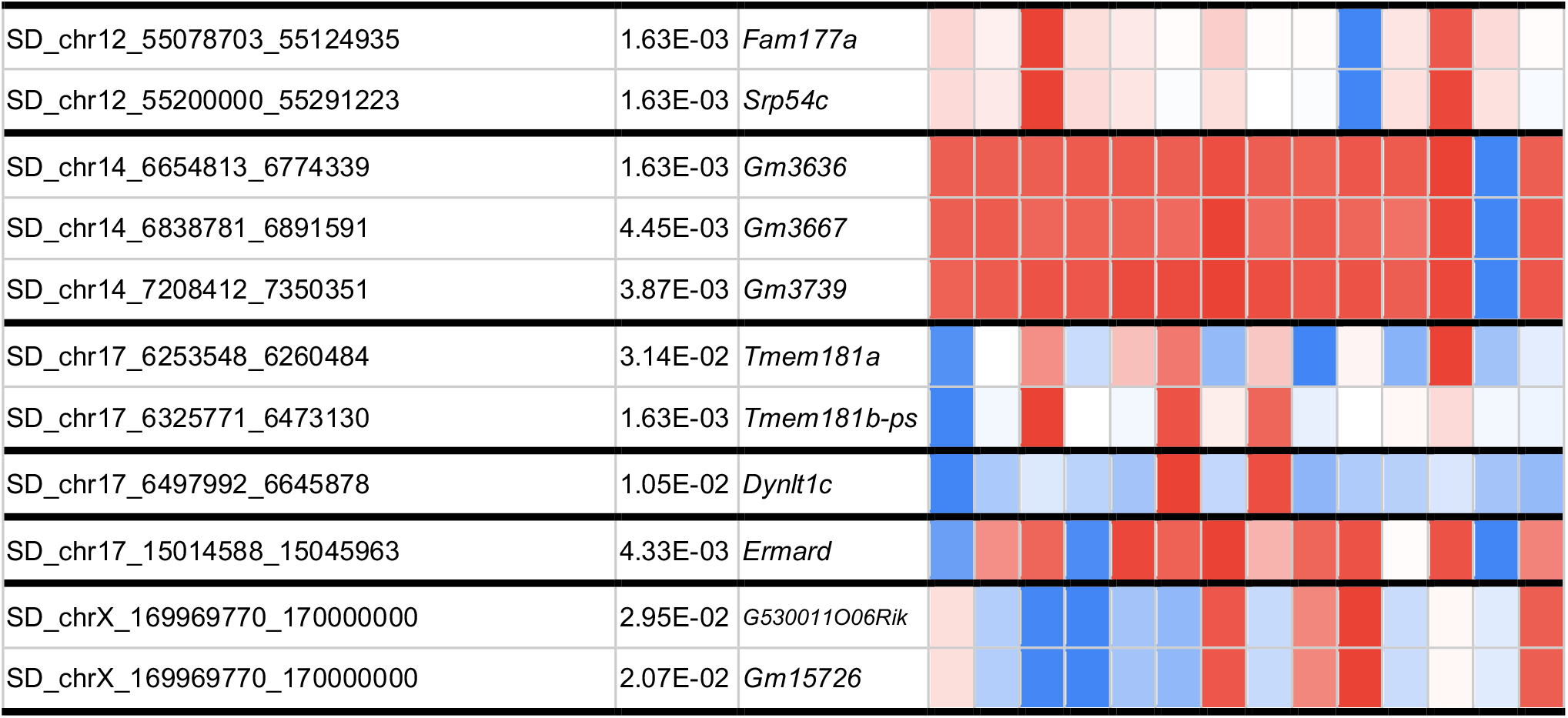
Significant associations and genotype patterns, related to Figure 2. Significant associations between DEGene expression and large effect variants with FDR<0.05 is presented. A linear mixed model is used with a Genomic Relatedness Matrix (GRM) to control for population structure as a random effect and parental strain (C57BL/6 versus C57BL/10) as a fixed effect to identify associations within C57BL/6 and C57BL/10 substrains. Variant information column indicates variant type (SD: segmental duplication), genomic coordinates and variant consequences (for SVs and INDELs). Colors represent genotype patterns for the variants. For bi-allelic variants (SVs and SNP/INDELs), red and blue colors represent two genotypes, while for multiallelic copy number variants the normalized read depth varies between 0 and 1 where 0: blue, 0.5: white, and 1: red represent three genotypes. The same genotype patterns are clustered together for chromosomes such as 2 and 4, which shows that nearby genes in these regions have been affected by the same copy number variation patterns. Bold horizontal lines segregate nearby variants with similar genotype patterns.

**Table S3. Validation rates of WGS SNPs and INDELs using RNA-Seq data, related to Methods details**. (excel file)

**Table S4. Sanger sequencing validation of loss-of-function SNPs and INDELs, related to Methods details**. (excel file)

