## Supplementary material for "Polymorphic SNPs, short tandem repeats and structural variants are responsible for differential gene expression across C57BL/6 and C57BL/10 substrains": Key resources table

| REAGENT or RESOURCE | SOURCE | IDENTIFIER |
|---|---|---|
| Experimental models: Cell lines | | |
| Mouse cell line: C57BL/6J *Upf2*-KO ES cells | Laboratory of Miles Wilkinson | N/A |
| Mouse cell line: C57BL/6J *Upf2*-WT ES cells | Laboratory of Miles Wilkinson | N/A |
| Experimental models: Organisms/strains | | |
| Mouse: C57BL/6NTac | Taconic | Model No: B6 |
| Mouse: C57BL/6NJ | The Jackson Laboratory | Stock No: 005304 |
| Mouse: C57BL/6NHsd | Harlan | Order code: 044 |
| Mouse: C57BL/6NCrl | Charles River | Strain code: 027 |
| Mouse: C57BL/6JEiJ | The Jackson Laboratory | Stock No: 000924 |
| Mouse: C57BL/6JBomTac | Taconic | Model No: B6JBom |
| Mouse: C57BL/6J | The Jackson Laboratory | Stock No: 000664 |
| Mouse: C57BL/6ByJ | The Jackson Laboratory | Stock No: 001139 |
| Mouse: B6N-TyrC/BrdCrCrl | Charles River | Strain code: 493 |
| Mouse: C57BL/10SnJ | The Jackson Laboratory | Stock No: 000666 |
| Mouse: C57BL/10ScSnJ | The Jackson Laboratory | Stock No: 000476 |
| Mouse: C57BL/10ScNHsd | Taconic | Model No: 046 |
| Mouse: C57BL/10ScCr | The Jackson Laboratory | Stock No: 003752 |
| Mouse: C57BL/10J | The Jackson Laboratory | Stock No: 000665 |
| Software and algorithms | | |
| HiSat2 v.2.1.0 | Kim et. al. 2019 | http://daehwankimlab.github.io/hisat2/ |
| HTSeq | Anders et. al. 2015 | https://htseq.readthedocs.io/en/master/overview.html |
| SpeedSeq | Chiang et. al. 2015 | https://github.com/hall-lab/speedseq |
| STAR v2.7.9a | Dobin et. al. 2013 | https://github.com/alexdobin/STAR |
| GATK v4.2.2.0 | Van der Auwera GA and O'Connor BD. 2020 | https://gatk.broadinstitute.org/hc/en-us |
| Plink v1.9 | Purcell et. al. 2007 | https://www.cog-genomics.org/plink/ |
| edgeR | Robinson et. al. 2010 | https://bioconductor.org/packages/release/bioc/html/edgeR.html |
| HipSTR v0.6 | Willems et. al. 2017 | https://hipstr-tool.github.io/HipSTR/ |
| LUMPY | Layer et. al. 2014 | https://github.com/arq5x/lumpy-sv |
| CNVnator | Abyzov et. al. 2011 | https://github.com/abyzovlab/CNVnator |
| mosdepth | Pedersen and Quinlan 2018 | https://github.com/brentp/mosdepth |
| Limix v3.0.4 | Lippert et. al. 2014 | https://horta-limix.readthedocs.io/en/api/installation.html |

| | | |
|---|---|---|
| Variant Effect Predictor (VEP) | McLaren et. al. 2016 | https://m.ensembl.org/info/docs/tools/vep/script/vep_download.html |
| sashimi_plot: used in Figure 3D | Katz et. al. 2010 | https://miso.readthedocs.io/en/fastmiso/sashimi.html |
| R v4.1.1 | R Core Team 2021 | https://www.r-project.org |
| RNAseq and WGS short read data | This paper | SRA: PRJNA705216 |
| Custom codes required to generate figures and perform analyses in this paper | This paper | https://zenodo.org/record/5716285 |
| Data required to run our custom codes and reproduce the figures and results | This paper | Mendeley DOI: 10.17632/39sw8xcrmv.2 |
| Processed data generated in this paper | This paper | Mendeley DOI: 10.17632/k6tkmm6m5h.3 |
